# Neurophysiology of effortful listening: Decoupling motivational modulation from task demands

**DOI:** 10.1101/2024.03.27.587038

**Authors:** Frauke Kraus, Bernhard Ross, Björn Herrmann, Jonas Obleser

**Author notes:** joint senior authorship (Björn Herrmann, Jonas Obleser). Correspondence concerning this article should be addressed to Björn Herrmann, Rotman Research Institute, Baycrest, 3560 14 Bathurst St, North York, ON, M6A 2E1, Canada, or to Jonas Obleser, Department of Psychology, University of Lübeck, Maria-Goeppert-Str. 9a, 23562 Lübeck, Germany,.

## Abstract

In demanding listening situations, a listener’s motivational state may affect their cognitive investment. Here, we aim to delineate how domain-specific sensory processing, domain-general neural alpha power, and pupil size as a proxy for cognitive investment encode influences of motivational state under demanding listening. Participants performed an auditory gap-detection task while pupil size and the magnetoencephalogram (MEG) were simultaneously recorded. Task demand and a listener’s motivational state were orthogonally manipulated through changes in gap duration and monetary-reward prospect, respectively. Whereas task difficulty impaired performance, reward prospect enhanced it. Pupil size reliably indicated the modulatory impact of an individual’s motivational state. At the neural level, the motivational state did not affect auditory sensory processing directly but impacted attentional post-processing of an auditory event as reflected in the late evoked-response field and alpha power change. Both pre-gap pupil dilation and higher parietal alpha power predicted better performance at the single-trial level. The current data support a framework wherein the motivational state acts as an attentional top-down neural means of post-processing the auditory input in challenging listening situations.

**Significance Statement:** How does an individual’s motivational state affect cognitive investment during effortful listening? In this simultaneous pupillometry and MEG study, participants performed an auditory gap-detection task while their motivational state was manipulated through varying prospect of a monetary reward. The pupil size directly mirrored this motivational modulation of the listening demand. The individual’s motivational state also enhanced top-down attentional post-processing of the auditory event but did neither change auditory sensory processing nor pre-gap parietal alpha power. These data suggest that a listener’s motivational state acts as a late attentional top-down effect on auditory neural processes in challenging listening situations.

## Introduction

The motivational state of a person influences whether they are willing to invest cognitively in demanding listening contexts (Brehm & Self, 1989; Kraus et al., 2023a; Richter et al., 2016). That is, a person who sees no intrinsic (e.g., enjoyment) or extrinsic (e.g., financial) value in listening, and is thus in a low motivational state, may be more likely to disengage from listening than someone who is highly motivated, especially under highly demanding conditions (Brehm & Self, 1989; Herrmann & Johnsrude, 2020; Pichora-Fuller et al., 2016; Richter et al., 2016). When listening conditions are less demanding, a person may still engage in listening despite being little motivated (Brehm & Self, 1989; Richter et al., 2016). Although this interacting influence of task demands and motivation (see Fig. 1C) is established theoretically and empirically, for example, for cognitive control tasks (Parro et al., 2018; Yee & Braver, 2018), little research has been conducted to investigate the impact of task demands and motivation on listening under challenges and to characterize the underlying neurophysiological mechanisms.

**Figure 1.**
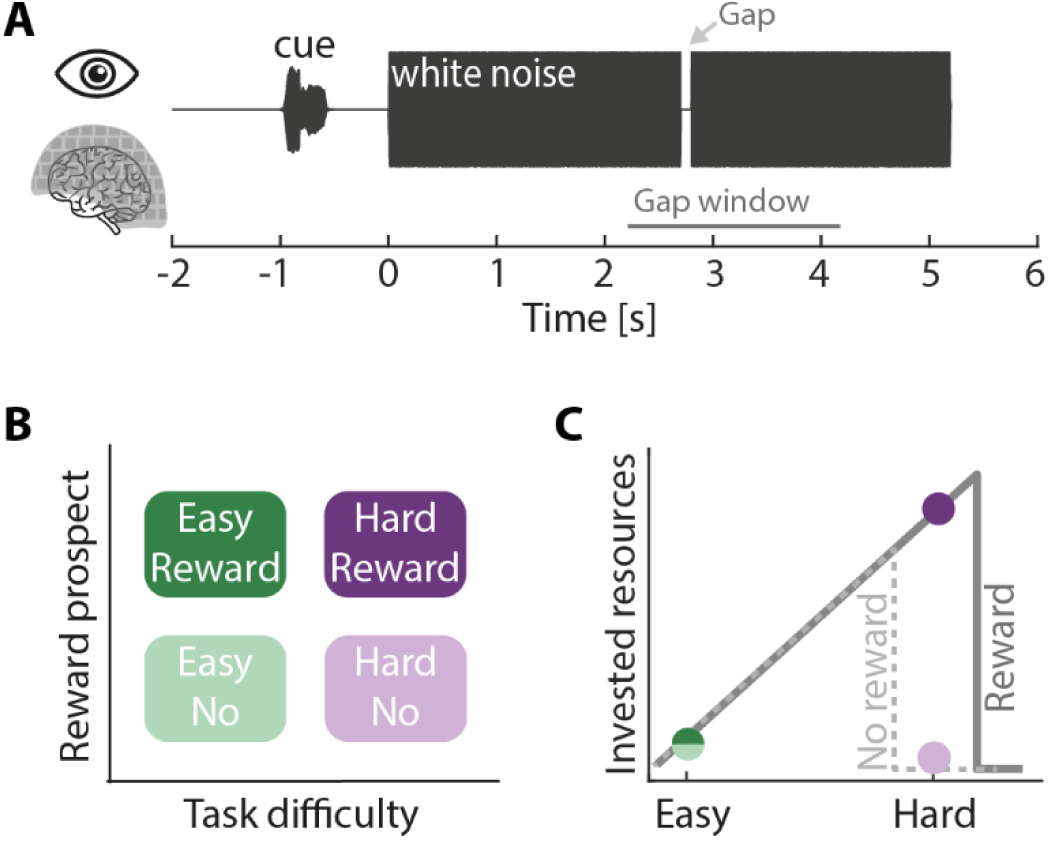
Experimental design. **A:** Auditory gap-detection task: Participants’ task was to detect a gap within 5.2 s white-noise sound. The gap occurred randomly between 2.2–4.2 s post noise onset (gap window, marked by the gray line). **B**: Two-by-two design: Task difficulty was determined by the gap duration (titrated to 65% detection performance for the hard condition and twice as long in easy trials). An auditory cue 1 s before each trial indicated whether the upcoming trial was reward-relevant or not. **C**: Hypothesis: According to the Motivation Intensity Theory (gray lines; Brehm & Self, 1989; Richter, 2016), participants should invest cognitively under the hard listening condition only when motivated to succeed (solid line), but should give up investing resources when they are less motivated (dashed line). The colored dots show our hypothesis. Figure design follows Kraus et al., (2023a).

**Figure 2.**
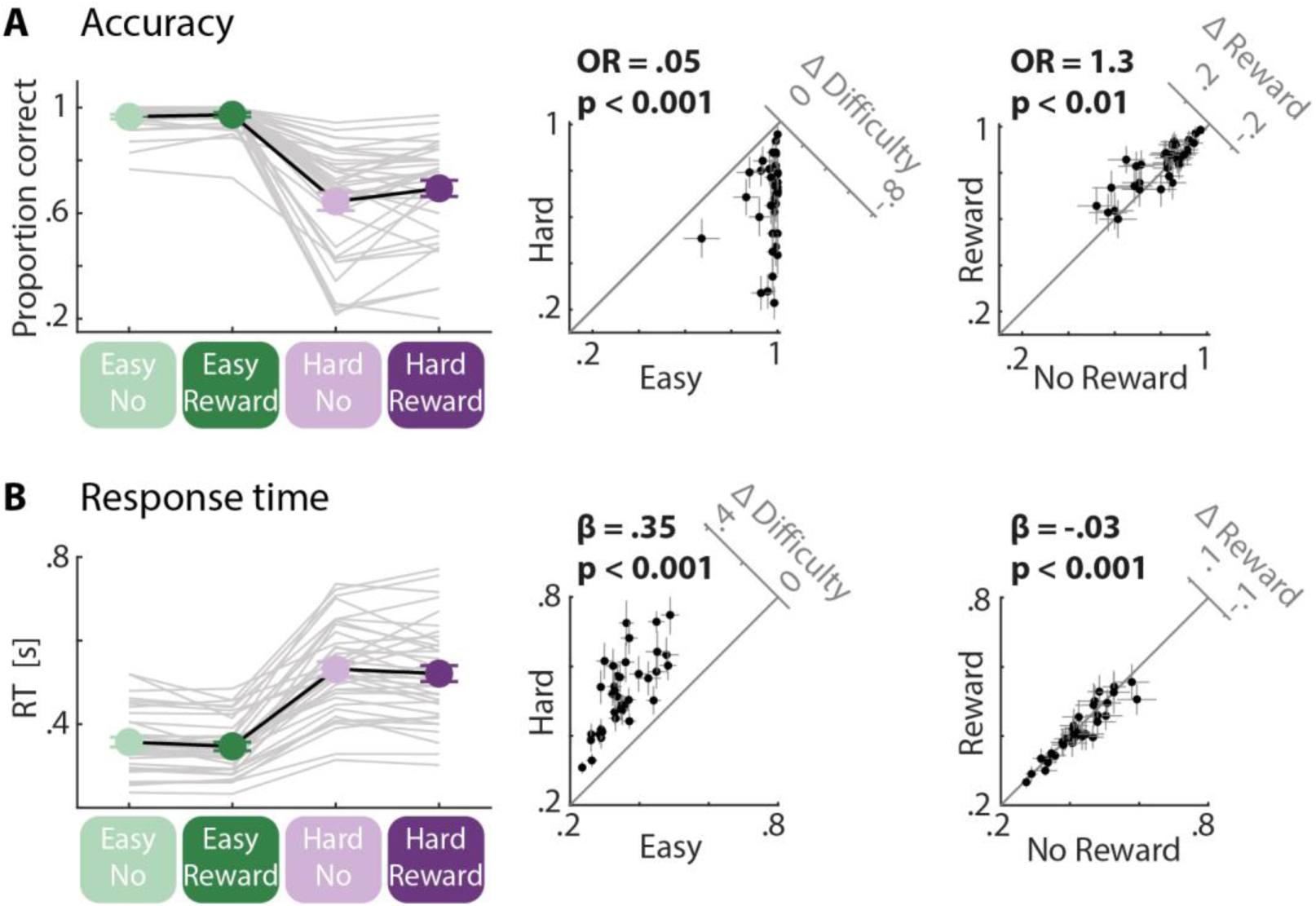
Behavioral results. **A:** Accuracy. Proportion correct was better for the easy compared to the hard condition and for the reward-relevant compared to the reward-irrelevant condition. Insets: 45-degree scatter plots showing the task difficulty (left) and the reward prospect effect (right) from linear mixed-model analysis. Difference plots (y minus x axis) are shown in the upper right corners). **B**: Response time. Participants were faster for the easy compared to the hard condition and for the reward-relevant compared to the reward-irrelevant condition. Insets: 45-degree scatter plots showing the task difficulty (left) and the reward prospect effect (right) from linear mixed-model analysis. Difference plots (y minus x axis) are shown in the upper right corners).

The noradrenergic (NE) modulation emerging from locus coeruleus (LC) in the brainstem is one driver of pupil dilation (Joshi et al., 2016; Joshi & Gold, 2020). The LC–NE system is sensitive to attention (Vazey et al., 2018) and has a role in optimizing task performance and task engagement (Aston-Jones & Cohen, 2005a). Accordingly, pupil dilation as a proxy of LC–NE activity has often been used as an indicator of cognitive investment: The pupil dilates as a task becomes more difficult (Kadem et al., 2020; Kahneman & Beatty, 1966; Koelewijn et al., 2012; Ohlenforst et al., 2018; Wendt et al., 2016; Winn et al., 2015; Zekveld et al., 2010). Moreover, pupil size dynamics during high listening demands are known to be modulated by an individual’s motivational state (Alfandari et al., 2023; Bijleveld et al., 2009; Zhang et al., 2019); recently confirmed by us using highly-controlled noise stimuli (Kraus et al., 2023a) that will also be used the present study. The motivation-driven additional cognitive investment indexed by pupil dilation predicted better behavioral outcomes (Kraus et al., 2023a).

Neural oscillatory alpha power is sensitive to listening demands, likely reflecting top-down attentional modulation (Dimitrijevic et al., 2017; Herrmann et al., 2023; Kraus et al., 2023b; Obleser et al., 2012; Paul et al., 2021; Petersen et al., 2015; Weisz et al., 2011; Wöstmann et al., 2015). However, in studies using highly controlled stimulus designs, in which task difficulty was manipulated without changing stimulus acoustics, an increase in alpha power has only been observed sometimes, for example, when relevant sound information occurs shortly after stimulus onset (Herrmann et al., 2023), but not when relevant sound information occurs late in a stimulus and when visual stimuli are presented concurrently (Kraus et al., 2023b). Manipulating motivation orthogonally to task demands may help identifying the neural correlates of cognitive investment during listening. That is, neural activity in brain regions that mediate top-down control of attention (e.g., cingulate cortex) is modulated by motivation (Small et al., 2005). Consistently, occipital alpha power in a visual search task has been sensitive to monetary rewards (Sawaki et al., 2015), raising the possibility that alpha power may be modulated by a person’s motivational state during listening, and, in turn, providing an opportunity to identify the neural mechanisms underlying challenging listening.

Apart from neural signals possibly indexing attention networks directly, neural activity elicited by auditory stimuli may provide additional insights into motivation influences on neural processing during listening. Auditory sensory responses around 100 ms after stimulus onset are enhanced as the saliency of a sensory input increases (for a review: Näätänen & Picton, 1987) and when individuals attend to, compared to ignore, auditory stimuli (Hillyard et al., 1973) but effects of motivation have not been observed (Goldstein et al., 2006; Krebs et al., 2013). Motivation rather seems to modulate neural signals about 300 ms after stimulus onset (Baines et al., 2011; Goldstein et al., 2006; van den Berg et al., 2014).

The current study will use magnetoencephalography (MEG) to investigate the neurophysiological changes associated with listening demands under different motivational states (using a reward manipulation). We will use pupil-size data as a benchmark for modulatory influences of motivation on cognitive investment during listening (Kraus et al., 2023a). Using MEG, we will investigate whether reward prospect impacts auditory sensory processing and how reward prospect influences attentional processing under varying task demands (alpha power). In detail, we expect increased attentional processing under high motivation when listening is hard, but no additional motivational boast when listening is easy (see Fig. 1C). Additionally, we will investigate the relationship among changes in pupil size, sensory responses, and neural alpha power and to behavioral listening outcome.

## Methods

### Participants

Thirty-seven adults aged between 18 and 34 participated in the current study (mean = 23.43 years; SD = 4.05 years; 11 males and 26 females; all right-handed). None of the participants reported having a history of neural disorders or hearing problems. Participants gave written informed consent before participation and received an honorarium of $15 CAD per hour. Their motivation was manipulated through financial rewards. Participants could earn an additional $15 CAD depending on their behavioral performance. The study was conducted at the Rotman Research Institute at the Baycrest Academy for Research and Education in Toronto, Ontario, Canada. The study protocols are in accordance with the Declaration of Helsinki and were approved by the local ethics committee of the Rotman Research Institute at the Baycrest Academy for Research and Education in Toronto, Ontario, Canada.

### Experimental environment

MEG data were recorded with a 275-channel axial gradiometer MEG device (CTF-MEG, Coquitlam, BC, Canada) in an electromagnetically shielded room. Participants were placed in a seating position with their head centered in the MEG helmet. A back-projection screen was placed about 70 cm in front of participants. An eye-tracking camera (Eyelink 1000 Plus, SR Research) was mounted to the screen. The experimental stimulation was controlled by a laptop (Windows 10) running Psychtoolbox (version 3.0.14) in MATLAB. Sounds were presented via an external sound card (RME Fireface UCx ii) and delivered binaurally via MEG-compatible insert earphones (EARTONE 3A, 3M). A button response box was used to record participants’ behavioral responses. After the MEG experiment, the location of the three fiducials (left tragus, right tragus, nasion) and the shape of the head were digitized using a 3-D digitizer (Polhemus Fastrak).

### Experimental design

The experimental design was similar to the one used in Kraus et al. (2023a). An auditory gap-detection task was chosen because the timing and degree of listening demand can be more tightly controlled than for speech materials that have been used in previous studies (Kraus et al., 2023a). In fact, our previous work shows effects of motivation on behavior and pupil size, and a critical interaction in line with theoretical frameworks (Brehm & Self, 1989; Richter et al., 2016; Figure 1C), which studies using speech materials have not been able to observe. The experimental control afforded by the auditory gap-detection task (see also Henry et al., 2014, 2017; Henry & Obleser, 2012; Herrmann et al., 2023; Kraus et al., 2023a, 2023b) facilitates the analysis of the interaction of motivation, listening demand and listening outcome (Kraus et al., 2023a).

Participants were presented with 5.2-s white-noise sounds that each contained one gap (Fig. 1) and pressed a button with the right index finger on the response box as soon as they detected the gap. The gap could occur at one of 70 randomly selected time points between 2.2 and 4.2 s after noise onset (linearly spaced). Auditory stimuli were presented at 45 dB above the participant’s sensation level, which was determined for each ear prior to the experiment using an audiometer. There was no visual stimulation during the presentation of the white noise. Participants were asked to keep their gaze on the black screen and not close their eyes.

The 2 × 2 design included the factors of task difficulty (easy, hard) and reward prospect (reward-irrelevant, reward-relevant). Task difficulty was manipulated via the duration of the gap in the noise sound. In the hard condition, gap duration was individually titrated to about 65% gap-detection performance in training blocks prior to the main experimental blocks (4–6 training blocks of 2 min each). Gaps in the easy condition were twice as long (Kraus et al., 2023a, 2023b).

Half of the trials were paired with the possibility of receiving an additional financial reward based on the participant’s performance on these trials (reward-relevant trials). In contrast, the other half of the trials were not paired with any reward (reward-irrelevant trials). Specifically, after the experiment, three trials for each of the two difficulty levels were chosen from the reward-relevant trials (Cole et al., 2022; Kraus et al., 2023a; Teoh et al., 2020; Tusche & Hutcherson, 2018). Participants could gain $15 CAD in addition to their hourly compensation rate if their average performance across these six trials was above 80%. All of the participants reached this threshold and got the reward.

Trials were presented in 14 blocks, each containing 20 trials. In half of the blocks, task difficulty was easy, whereas task difficulty was hard in the other half. Easy and hard blocks alternated. The difficulty of the starting task was counterbalanced across participants. At the beginning of each block, participants received written information about the task difficulty (easy or hard) of the upcoming block. Hence, participants had prior knowledge about whether detecting the gap would be easy or hard. In random order, half of the 20 trials per block were reward-relevant trials, whereas the other half were reward-irrelevant trials.

An auditory cue consisting of a guitar or a flute sound occurred before each white noise sound, indicating whether a trial was reward-relevant or reward-irrelevant. The pairings of guitar and flute sound to reward-relevant and reward-irrelevant trials were counterbalanced across participants. Participants familiarized themselves with the cue-reward association during training trials before the experiment. Overall, the experiment contained 70 white-noise sounds per Task difficulty (easy, hard) × Reward prospect (reward-irrelevant, reward-relevant) condition, resulting in 280 trials.

After the experiment, participants completed a questionnaire regarding their use of the reward cues. They first indicated which auditory cue was associated with reward-relevant trials. We then asked them to rate the following statement: “I used the auditory cues (guitar and flute) to distinguish between important and unimportant conditions” on a 6-point scale ranging from “strongly disagree” over “disagree”, “somewhat disagree”, “somewhat agree”, “agree” to “strongly agree”. Participants rated this statement twice, separately for the easy and the hard conditions.

### Analysis of behavioral data

Any button press within 0.1 to 1.5 s after gap onset was defined as a hit (coded 1). Trials for which no button was pressed within this time window were considered a miss (coded 0). The time between the gap onset and the button press was calculated as the response time. The time at which the gap occurred within the sound was included in the statistical modeling (see Statistical Analysis) to account and test for an expected hazard effect (higher accuracy and faster response times for later compared to early gaps relative to white-noise onset; Herrmann et al., 2023; Niemi & Näätänen, 1981; Nobre et al., 2007). Response-time data were log-transformed for statistical analysis to obtain values closer to a normal distribution.

### Pupil data recording and preprocessing

Eye movements and pupil size of the right eye were continuously recorded using an Eyelink 1000 Plus eye tracker (SR research). Data was recorded as an additional channel of the MEG data acquisition. MATLAB (MathWorks, Inc.) was used for the preprocessing and analyses of the data. To detect eye blinks, we determined the threshold in the pupil channel above which the eye-tracker had lost the pupil in case of a blink. All datapoints above this threshold were coded as invalid (NaN). Additionally, 100 ms before and 200 ms after above threshold time points were also coded as NaN. NaN-coded data points were linearly interpolated. Pupil data were subsequently low-pass filtered at 4 Hz (Butterworth, 4^th^ order) and divided into epochs ranging from -2 to 6.2 s time-locked to white-noise onset. An epoch was excluded if more than 40% of the trial had been NaN-coded prior to interpolation. If more than 50% of the trials in any condition were excluded, the full pupil dataset of the respective participant was excluded from analysis (N=12). The high number of excluded pupil datasets is in part due to the technical challenge combining the concurrent recording of pupil size and MEG. Pupil-size data were downsampled to 50 Hz. The polarity of the pupil data was inverted at the analysis stage because the pupil recordings through the MEG were inverted (we determined this in a brief black vs. white screen experiment to elicit dilation vs constriction, respectively). For each trial, the mean pupil size in the -1.5 to -1.1 s time window was subtracted from the pupil size at every time point of the epoch (baseline correction). This baseline time window was chosen to avoid contamination of the auditory cue which was presented at -1 s. For each participant, single-trial time courses were averaged, separately for each condition.

### Pupil size analysis

For the statistical analysis (described in the Statistical analysis section) of the effects of task difficulty and reward prospect, pupil-size data were averaged across the time window ranging from 2.2 s (onset of gap window) to 6.2 s (end of trial).

Similar to our previous approach (Kraus et al., 2023a), we investigated whether a smaller pupil size is associated with an increased probability that a participant misses a gap or responds slower to the gap. Therefore, pupil-size data (baseline-corrected to -1.5 to -1.1 s to noise-onset) were time-locked to the gap onset. For statistical analysis, pupil size was averaged across the -.5 to 0 s time window, time-locked to gap onset.

To illustrate the association between pupil size and response time (Fig. 3E), we created two groups of trials: fast and slow trials. Fast trials were defined as those with response times equal to or faster than 0.75 SD of the mean. Slow trials were defined as those with response times equal to or slower than 0.75 SD of the mean. The threshold was chosen as a good compromise between too many and too few trials per group. The grouping was done separately for reward-relevant and reward-irrelevant trials to ensure the same number of trials were used for each group and reward condition. Additionally, we included only trials into the group of fast trials if their gap-onset time matched with one trial of the slow trials and vice versa to ensure that there was no difference in gap-onset times between slow and fast trials. The pupil size of the averaged slow versus fast trials was compared using a one-sample t-test per time point. To account for multiple comparisons, p-values were corrected across time points using a false discovery rate (FDR) of q = 5% (Benjamini & Hochberg, 1995).

**Figure 3.**
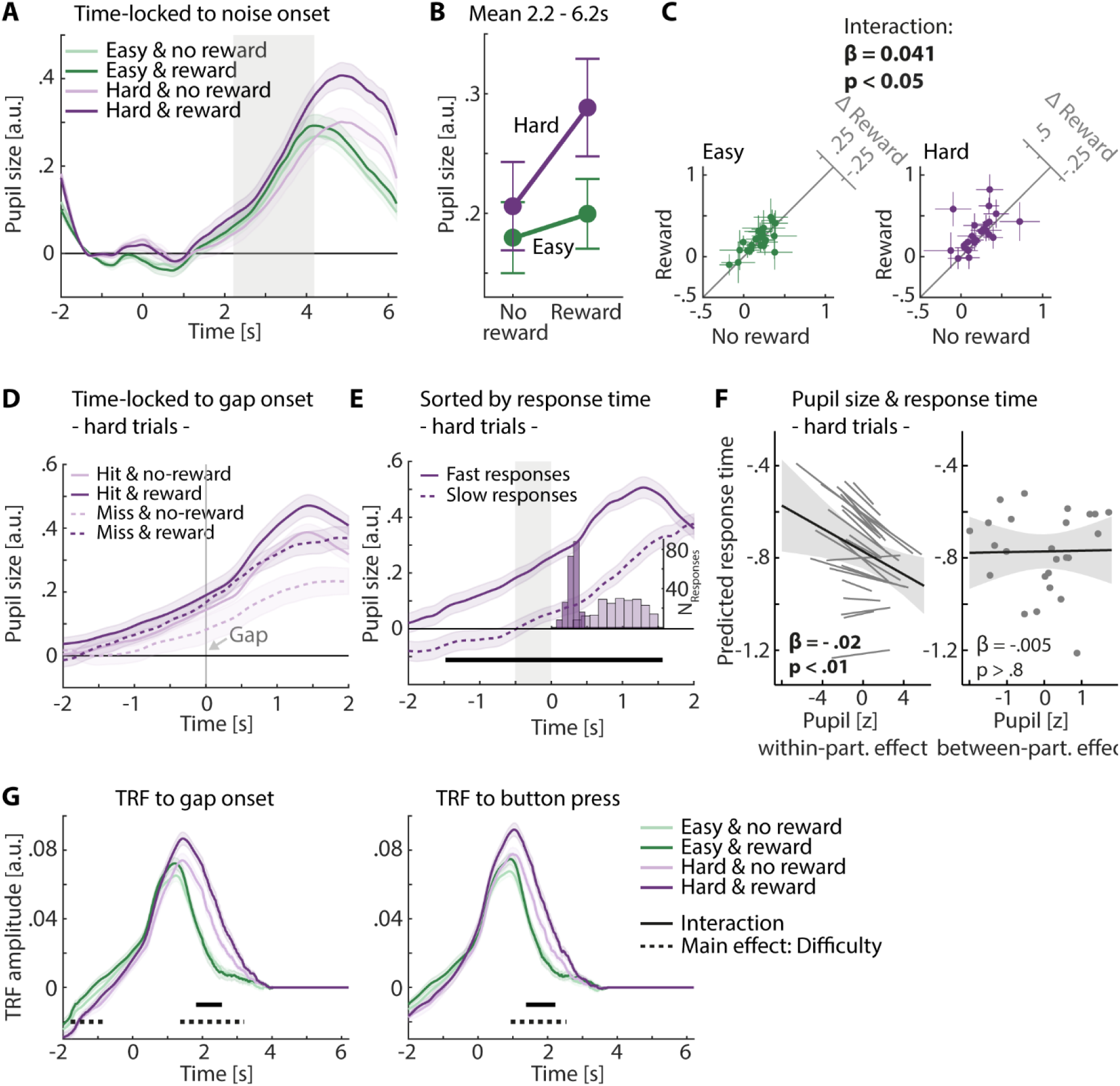
Pupil size results. **A:** Averaged pupil-size time courses across participants per condition. Error bands reflect the within-participants error. Gray areas indicate time window during which a gap could occur, from 2.2 to 4.2 s. **B**: Averaged data for 2.2–6.2 s time window. Error bars indicate the standard error of the mean. **C**: 45-degree scatter plots illustrate the interaction. Left: Data from the easy condition. Right: Data from the hard condition. Colored dots show averaged pupil data per task-difficulty level, separately for each participant. The 45-degree line indicates no difference between conditions. Crosshairs indicate the 95% confidence interval (CI). Difference plots (y-minus x-axis) are shown in upper right corners. **D**: Pupil-size time courses (time-locked to gap onset) for the hard condition, separately for each reward-prospect condition (light vs. dark) and for hit and miss trials (solid vs. dashed lines). Error bands reflect the within-participants error. **E**: Pupil-size data grouped into trials with slow (dashed) and fast response times (solid). The histogram shows the distribution of response times for each group. Gray area indicates the time window for the linear mixed-effect model analysis in panel F. Black line indicates the time window in which the pupil size was significantly larger for fast-response trials compared to slow-response trials (after FDR-correction). Error bands reflect the within-participants error. **F**: Effect of pupil size on response time in a linear mixed-effect model analysis. A larger pupil size is associated with a faster response time on a within-participant level (not for the between-participants effect). Participant-specific slope for pupil-size did not improve the model, but we show it here for illustrative purposes. **G:** TRF-approach of the pupil-size response to the gap onset (left) and button press (right). Black line indicates the time window in which the pupil size increased stronger for reward-relevant than reward-irrelevant trials when task difficulty was hard compared to easy. Dashed black line indicates the time window in which the pupil size was significantly larger in hard compared to easy trials. All shown significant effects are based on FDR-correction.

### MEG recording and preprocessing

Magnetoencephalography (MEG) data were recorded using a CTF-MEG (Coquitlam, BC, Canada) with 275 axial gradiometers at a sampling frequency of 1200 Hz.

MEG data analysis was performed with the Fieldtrip MATLAB toolbox (2019-09-20; Oostenveld et al., 2011). To avoid numerical issues during processing due to very small numbers associated with MEG recordings in the Tesla range, we multiplied all MEG data with 10^12^, leading to data in the picotesla range. Data were filtered with a 100 Hz low-pass (Hann window, 89 points), a 0.7 Hz high-pass (Hann window, 2869 points), and a 60 Hz elliptic band-stop filter to suppress line noise. Data were cut into 10.2-s trials ranging from -3 to 7.2 s time-locked to noise onset. For the calculation of independent component analysis (ICA), these trials were divided into 1-s snippets. Components containing blinks, lateral eye movement, and heart-related activity were identified through visual inspection. The continuous data (filtered) were projected to ICA space using the unmixing matrix. The previously identified components containing artifacts were removed from the continuous data and the data were then back-projected to the original 275 sensors using the mixing matrix. Data were then low-pass filtered at 30 Hz (Hann window, 111 points) and divided into trials of 10.2 s (−3 to 7.2 s time-locked to noise onset). Data were downsampled to 600 Hz, and trials that exceeded a signal change of more than 4 picotesla in any of the MEG channels were excluded from analysis.

### Analysis of event-related fields for gap onset and button response

Data were transformed from axial to planar gradiometers (Vrba & Robinson, 2001). Planar gradiometers show the strongest sensitivity to sources that originate from directly below them (Hämäläinen, 1995; Vrba & Robinson, 2001), which makes the interpretation of topographical distributions more intuitive. The signal at the two planar gradiometers forming one pair were combined by calculating the sum. This resulted in signals at 275 sensors.

For the analysis of event-related fields (ERFs), we focused on hit trials and each trial was time-locked to the respective gap-onset time. To investigate evoked auditory sensory processing, data were averaged across temporal sensors (MRF67, MRT13, MRT14, MRT15, MRT23, MRT24, MRT25, MRT2, MRT35, MRT36, MLF67, MLT13, MLT14, MLT15, MLT23, MLT24, MLT25, MLT26, MLT35, MLT36). For illustrative purposes, trials were averaged separately for each of the four conditions and baseline-corrected by subtracting the mean amplitude in the time window before gap onset (−.5 to 0 s) from each time point. For the statistical analysis, data were averaged across the time window of the M100-component of the ERF (0.09 to 0.13 s; Näätänen & Picton, 1987). The M100 time window was selected to investigate possible reward-prospect influences on auditory sensory processing.

For an exploratory analysis of the late ERF-component with a parietal topography (Figure 4A & C), hit trials were time-locked to a person’s response time in the respective trial and averaged across parietal sensors (MLC16, MLC17, MLC24, MLC25, MLC31, MLC32, MLC41, MLC42, MLC53, MLC54, MLC55, MLC61, MLC62, MLC63, MRC16, MRC17, MRC24, MRC25, MRC31, MRC32, MRC41, MRC42, MRC53, MRC54, MRC55, MRC61, MRC62, MRC63, MLP11, MLP12, MLP21, MLP22, MLP23, MLP31, MLP32, MLP33, MLP34, MLP35, MLP41, MLP42, MLP43, MLP44, MLP45, MLP54, MLP55, MLP56, MLP57, MRP11, MRP12, MRP21, MRP22, MRP23, MRP31, MRP32, MRP33, MRP34, MRP35, MRP41, MRP42, MRP43, MRP44, MRP45, MRP54, MRP55, MRP56, MRP57, MZC03, MZC04, MZP01).

**Figure 4.**
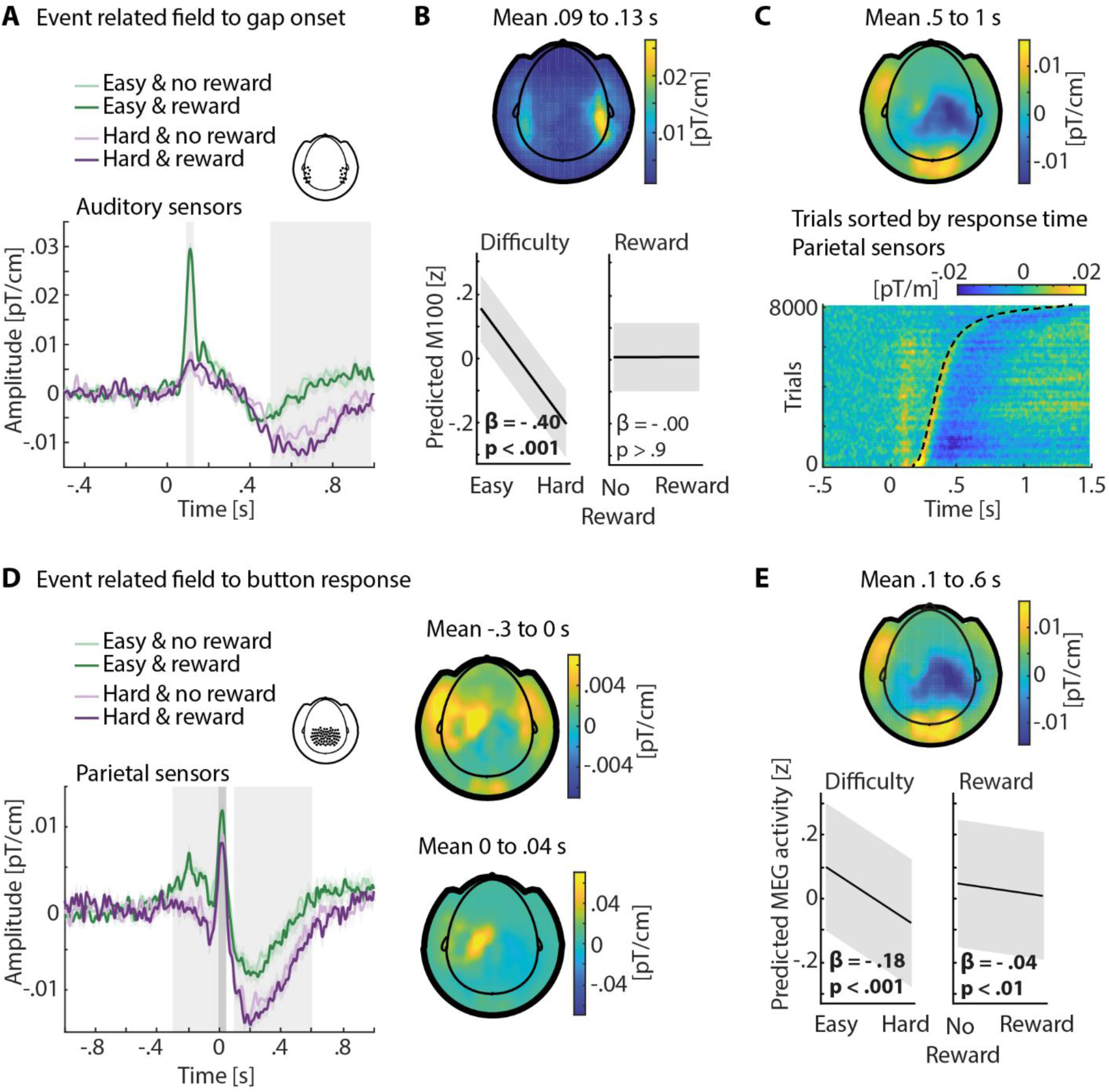
Event-related field to gap onset and to button response. **A:** Event-related field to gap onset. Data are averaged across temporal sensors marked in inset. Error bands reflect the within-participants error. Gray areas indicate time windows for analysis in B and C. **B**: Effect on M100. Average from .09 to .13 s. Top: Topography for the average across all for conditions. Bottom: Statistical results are calculated using a linear mixed-effect model. Main effect is significant for difficulty but not for reward prospect. **C**: Late ERF-component. Top: Topography for the average across all four conditions from .5 to 1 s. Bottom: Trials sorted by response time and time-locked to the gap-onset. Average across parietal sensors that are marked in the inset in panel D. Data are from all conditions. Black dashed line indicates response time. **D:** Event-related field to button response. Data are averaged across parietal sensors marked in the inset. Error bands reflect the within-participants error. Gray areas indicate time windows for topographies on the right, and in panel E. Topographies for time window before button response and around button response show main activity around left motor areas. **E:** After-response-component. Top: Topography for the average across all four conditions from .1 to .6 s. Bottom: Statistical analysis of time window .1 to .6 s after button response using a linear mixed-effect model. Stronger deflection for hard compared to easy trials and for reward-relevant compared to reward-irrelevant trials. All significant effects are based on FDR-correction.

For illustrative purposes, data were baseline-corrected using the same time window as for the gap-locked data (−.5 to 0 s time-locked to gap onset). For statistical analysis, single-trial response-locked data were averaged across the 0.1 to 0.6 s time window (time-locked to button response) to investigate task difficulty and reward-prospect effects on the response-related component (see Fig. 4D).

### Analysis of time-frequency power

In order to analyze oscillatory activity in the alpha frequency range (8 to 12 Hz; Jensen & Mazaheri, 2010; Klimesch et al., 2007; Weisz et al., 2011), single-trial time-domain data (after planar gradiometer calculation) were convolved with Morlet wavelets. Complex wavelet coefficients were calculated for frequencies ranging from 8–12 Hz (1 Hz steps) and time points ranging from –2 s to 6.2 s (0.04 s steps) time-locked to noise onset, separately for each trial, sensor, and participant. For the visualization of time-frequency data across a wider frequency range, calculations were also done for frequencies ranging from 1–20 Hz in steps of 1 Hz and in time steps of 0.16 s. We analyzed data first time-locked to noise onset and then time-locked to the gap onset.

For the analysis of data time-locked to noise onset (Fig. 5), power was calculated by squaring the magnitude of the complex wavelet coefficients separately for each trial, sensor, and time-frequency bin. Data for the two corresponding planar gradiometers of a pair were combined by calculating the sum, which resulted in power data for 275 sensors. For visualization purposes, time-frequency power per condition was averaged across trials, and baseline-corrected to dB power change: Data at each time point were divided by the mean power in the baseline time window (−1.5 to -1.1 time-locked to noise onset), and subsequently log10 transformed. To visualize the difference in power between hit and miss trials, trials were categorized as hit and miss trials and averaged across parietal sensors. For statistical analysis, parietal alpha power in the 1.7 to 2.2 s time window (time-locked to noise onset) was averaged. This time window was chosen because it precedes the time window during which a gap could occur (2.2–4.2 s).

**Figure 5.**
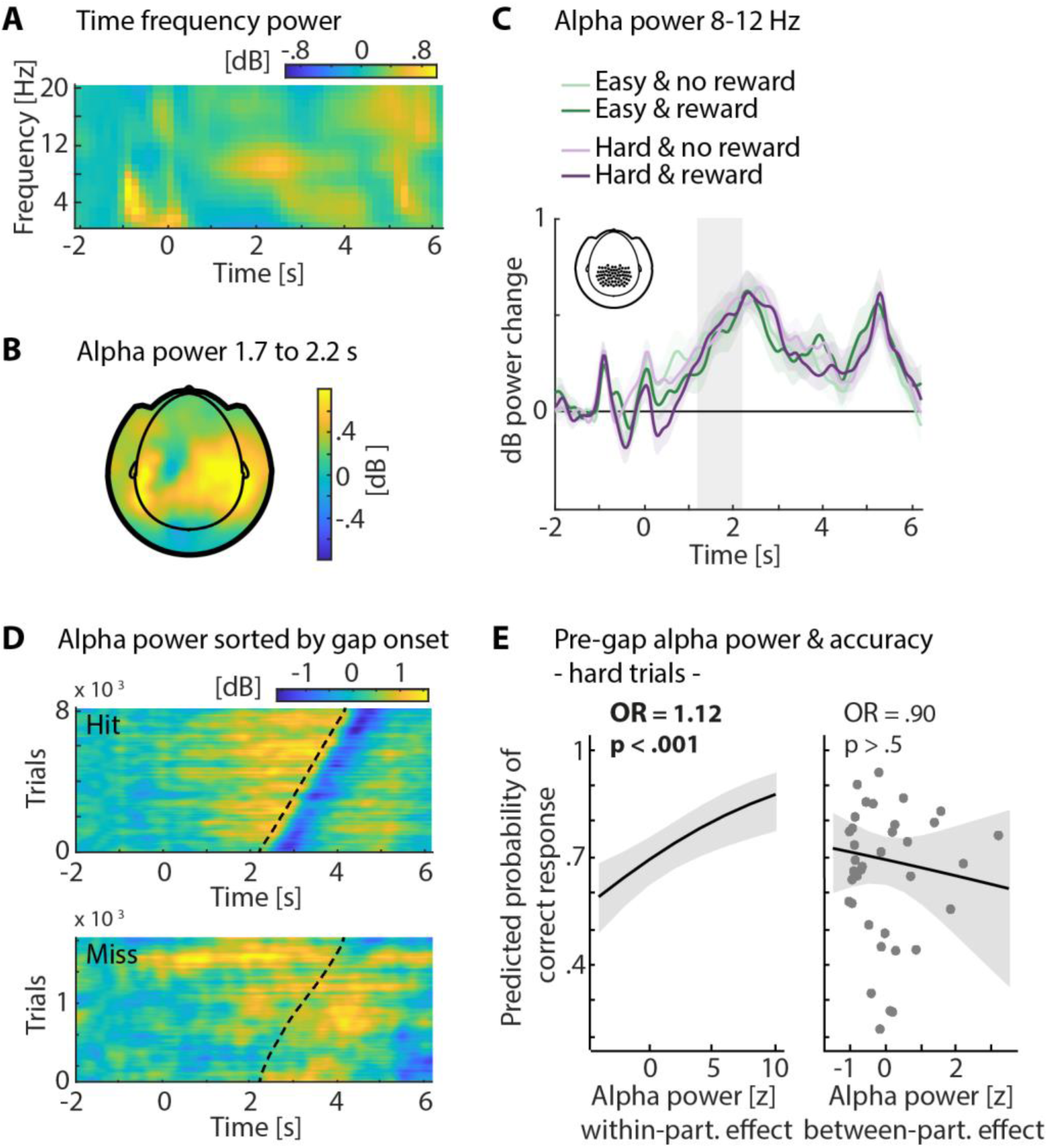
Pre-gap alpha power analysis. All data are averaged across parietal sensors marked in the inset in panel C. **A**: Time-frequency power measured in dB relative to a pre-stimulus time interval and averaged across all conditions. **B:** Topography reflects alpha power in the 500 ms (1.7-2.2 s) before the time window when a gap could occur (averaged across all conditions; gray area in C). **C:** Alpha power averaged for all four conditions separately. Error bands reflect the within-participants error. Effects of task difficulty and reward prospect were not significant in the 500 ms before a gap could occur (gray area). **D:** Alpha power for hit trials (top) and miss trials (bottom) sorted by gap onset. Black dashed lines indicate the gap time. Alpha power increased towards the gap but was suppressed after gap occurrence (for hit trials). **E:** Effect of pre-gap parietal alpha power on accuracy in a linear mixed-effect model analysis. Larger pre-gap alpha power is associated with better performance on a within-participant level only. Reported effects are based on FDR correction.

For the investigation of alpha activity changes related to gap detection, we time-locked the data to the gap onset and focused on hit trials only. This was done on single-trial complex wavelet coefficients, and power calculation and gradiometer combination were performed in the same way as for the noise-locked data above. For visualization purposes, condition-averaged data were baseline-corrected to dB power change using the same time window as for the noise-locked data (−1.5 to -1.1 s time-locked to noise onset) and separately averaged across parietal or temporal sensors (see section above). For the statistical analysis, pre-gap alpha power for each trial was averaged across the -0.5 to 0 s time window (time-locked to gap onset).

Trials in which participants detected the gap elicited an alpha-power suppression (Fig. 4 D). We thus also investigated possible effects of task difficulty and reward prospect on the post-gap alpha-power suppression. To this end, we calculated the latency of minimal alpha power between 0 and 1 s post-gap for each condition and participant. For the statistical analysis of the alpha power at its minimum, we averaged the data ±40 ms around the estimated latency for each trial.

### Source localization

For source localization, the MRI image of the standard brain and skull in Fieldtrip toolbox was non-linearly warped for each participant to fit their head-shape data from the Polhemus digitization. An automatic freesurfer’s approach was used to segment the MRI and extract the cortical mesh. This procedure ensured that the cortical mesh consisted of comparable grid-point locations across participants (Alavash et al., 2021) in accordance with the Human Connectome Project (HCP) standard atlas template (Glasser et al., 2017; Keitel & Gross, 2016). This cortical mesh served as the source model for each participant. Fieldtrip’s ‘singleshell’ method was used to calculate the inner skull surface that was used as a volume conductor model. Subsequently, leadfields were calculated separately for each experimental block to account for the slightly different head positions relative to the sensors in the different MEG recording blocks.

For each participant and block, a multi-taper (±2Hz spectral smoothing) fast Fourier transform (FFT) was calculated. The complex coefficients at 10 Hz that resulted from the FFT were used to calculate a cross-spectral density matrix per block. Dynamic imaging of coherent sources was used for source localization (DICS, Gross et al., 2001). For each source location, a spatial filter was calculated using a participant’s cross-spectral density matrix (dipole orientation: axis of most variance using singular value decomposition). For each trial, spatial filter weights were multiplied time point per time point with the complex wavelet coefficients from the analysis that used data time-locked to the gap onset (8–12 Hz frequency range). Power was calculated by squaring the magnitude of the complex wavelet coefficients that were projected through the spatial filter. Time-frequency source power was averaged across the 8 to 12 Hz frequencies.

For the statistical analysis, we focused on two predefined regions of interest (ROI) according to functional parcels defined by the Glasser atlas as available in the Human Connectome Project workspace (Glasser et al., 2017; Keitel & Gross, 2016). We included an auditory ROI due to the auditory nature of the task and because auditory stimulation is known to elicit sensory alpha activity (Herrmann et al., 2023; Kraus et al., 2023b; Mazaheri et al., 2014; Wöstmann et al., 2017). We also included a parietal ROI, because the parietal cortex is part of the attentional networks and is known to elicit alpha activity under task load (Banerjee et al., 2011; Behrmann et al., 2004; Herrmann et al., 2023; Kraus et al., 2023b; Rushworth et al., 2001). To analyze post-gap alpha-power suppression in source space, we calculated the latency of minimal alpha power between 0 and 1s post-gap per condition, participant, and ROI. For each trial, we then averaged the data ±40 ms around the estimated latency.

For visualization only, hit trials were averaged for each condition and participant. The dB power change was calculated relative to the original baseline time window (−1.5 to -1.1 s time-locked to noise onset).

### Statistical analysis

A paired sample t-test was used to analyze the subjective rating of cue use. The effect size was reported as Cohen’s d (J. Cohen, 1988).

A variety of linear mixed-effect models in R (v4.1.2), with the packages lme4 and sjPlot, were used for the statistical analyses. All linear mixed-effect models included participant-specific random intercepts to account for individual differences in the respective dependent variable. Single-trial accuracy data were binary. Hence, for accuracy data, a generalized linear mixed-effect model (GLMM) with a binomial distribution and a logit link function was used (Kraus et al., 2023a; Tune et al., 2021). All other models included continuous data as dependent variables, and we thus used a linear mixed-effect model (LMM) with a Gaussian distribution and an identity link function (Kraus et al., 2023a; Tune et al., 2021). All continuous variables were z-scored. All models predicting behavioral outcomes included the gap time as a regressor to account for hazard-rate effects (Herrmann et al., 2023; Niemi & Näätänen, 1981; Nobre et al., 2007). For all statistical models using neurophysiological data as dependent variable, data averaged across the baseline time window were included in the model as a separate regressor instead of using baseline-corrected data as the dependent variable (Alday, 2019).

First, linear mixed-effect models were calculated to analyze the influence of the two experimental factors on behavioral outcomes, pupil size, M100-component, and alpha power. The models included effects of task difficulty (easy, hard), reward prospect (reward-irrelevant, reward-relevant), and task difficulty × reward prospect interaction. Task difficulty and reward prospect were categorical predictors and were, therefore, deviation-coded (i.e., -0.5 [easy, no-reward] and 0.5 [hard, reward]).

Second, linear mixed-effect models were calculated to analyze the influence of pupil size or parietal alpha power on accuracy or response time. For these analyses, only hard trials were used because of the large variance in the behavioral measures between easy and hard conditions and the ceiling performance for the easy condition. Pupil size and parietal alpha power were used as independent variables in these models; therefore, baseline correction was needed before inclusion into the respective statistical model. Including the baseline data as an additional regressor (as described for the prediction of pupil/alpha data) would lead to collinearity issues. For pupil-size data, we used subtractive baselining (see Pupil size recording and preprocessing), which can be used on single trials (Mathôt et al., 2018). However, dB-power-baselining should not be used on single-trial data (M. X. Cohen, 2014). To account for this, we calculated a model with pre-gap alpha power as the dependent variable and alpha power within the baseline time window as the only independent variable. The residuals of this model served as the alpha power regressor in the models to predict behavioral outcomes (Alday, 2019).

To disentangle associations of pupil size/alpha power and behavior at the trial-by-trial state level (i.e., within-participant) from associations at the trait level (i.e., between-participants), we included two separate regressors associated with changes in pupil size/alpha power. The between-participants regressor contained trial-averaged pupil size/alpha power per individual, whereas the within-participant regressor contained the single-trial pupil size/alpha power relative to the individual mean (Bell et al., 2019; Kraus et al., 2023a, 2023b; Tune et al., 2021). Reward prospect and gap time were included as regressors to account for their potential influence on behavior. A random slope for the within-participant effect was included whenever it improved the respective model and did not lead to missing convergence of the model.

Third, a linear mixed-effect model was calculated to analyze the relationship between the M100 amplitude and the post-gap alpha power. Reward prospect, task difficulty, and their interaction were included as regressors to account for their potential influence on post-gap alpha power. A within-participant and a between-participants regressor were included for the M100-amplitude regressor as well as a random slope for the within-participant effect.

Fourth, linear mixed-effect models were calculated to analyze the relationship between the pre-gap pupil size and the M100-amplitude or the post-gap alpha power. Reward prospect, task difficulty and their interaction were included as regressors to account for their potential influence on the M100-amplitude or post-gap alpha power. A within-participant and a between-participants regressor were included for the pupil size regressor, as well as a random slope for the within-participant.

For the interpretation of effects analyzed with the different linear mixed-effect models, we calculated Bayes Factors (BF) as log(BF) = [BIC(H0)-BIC(H1)]/2, with BIC being the Bayes–Schwartz information criterion (Alavash & Obleser, 2024; Wagenmakers, 2007). To this end, we calculated the BIC for the full model, including the regressor of interest (H1), and for the reduced model, excluding the regressor of interest (H0). A log(BF) larger than 1 provides evidence for the presence of an effect of the regressor of interest, whereas a log(BF) value smaller than –1 suggests the absence of an effect of the regressor of interest (Dienes, 2014).

## Data availability

All data and analysis scripts are available at https://osf.io/prwb7/.

## Results

### Reward prospect improves performance

As intended, performance was better in the easy compared to the hard condition for both accuracy and response time (accuracy: GLMM; odds ratio (OR) = .05, std. error (SE) = .004, p = 1.74 × 10^-238^, log(BF) = Inf; response time: LMM; β = 0.35, SE = .007, p < 1.74 × 10^-238^, log(BF) = Inf; Fig. 2). Furthermore, participants were more accurate and faster for the reward-relevant compared to the reward-irrelevant condition (accuracy: OR = 1.31, SE = .117, p = 3.54 × 10^-3^, log(BF) = -.15; response time: β = -.03, SE = .007, p = 8.87 × 10^-5^, log(BF) = 3.35). The interaction was not significant (accuracy: odds ratio (OR) = 0.98, SE = .175, p > .9, log(BF) = -4.6; response time: β = -.02, SE = .014, p > .2, log(BF) = -3.75). We found a hazard-rate effect, such that individuals were more accurate and faster when the gap occurred later rather than earlier (accuracy: OR = 1.21, SE = .036, p = 3.19 × 10^-10^, log(BF) = 15.8; response time: β = -.08, SE = .003, p = 2.44 × 10^-118^, log(BF) = 263.5).

Participants rated their use of the auditory cue to distinguish between reward-relevant and reward-irrelevant higher in the hard compared to the easy condition (t_36_ =3.54, p = 1.11 × 10^-3^, d =.58).

### Reward prospect modulates the task difficulty effect on pupil size

To investigate the influence of task difficulty and reward prospect on pupil size, we averaged pupil size over the time window ranging from 2.2 to 6.2 s (Fig. 3B; time-locked to noise-onset). Pupil size was larger for hard compared to easy trials (β = .05, SE = .009, p = 5.21 × 10^-7^, log(BF) = 9.1) and larger for reward-relevant compared to reward-irrelevant trials (β = .03, SE = .009, p = 3.32 × 10^-4^, log(BF) = 2.6). Importantly, we observed a significant task difficulty × reward prospect interaction, indicating a larger increase in pupil size for reward-relevant than reward-irrelevant trials when task difficulty was hard compared to easy (β = .04, SE = .018, p = 3.08 × 10^-2^, log(BF) = -1.9; Fig. 3C).

Next, we investigated whether a larger pupil size during the interval preceding the gap is associated with behavioral performance. This analysis was conducted using the hard trials only and the information about the reward conditions was included as a regressor into the model. With this approach we controlled for the effects of reward prospect and task difficulty on both pupil size and response time. To this end, pupil-size data for hard trials were aligned to gap-onset times, and the pupil size was averaged across the -0.5 s to 0 s pre-gap time window. Pre-gap pupil size did not predict behavioral accuracy, neither at the trial-by-trial state level (within-participant: OR = 1.07, SE = .043, p > .1, log(BF) = -2.7) nor at the trait level (between-participants: OR = 1.04, SE = .158, p > .8, log(BF) = -1.6). However, at the trial-by-trial state level response times were shorter when pre-gap pupil size was larger (within-participant: β = -.02, SE = .008, p = 9.03 × 10^-3^, log(BF) = 0), but at the trait level there was no association between a participant’s mean pupil size and mean response time (between-participants: β = -.01, SE = .038, p > .8, log(BF) = -1.6; Fig. 3F). To illustrate this effect, we grouped the hard trials into fast and slow response trials, separately for the reward-relevant and the reward-irrelevant condition (trials were selected such that the mean gap time for fast and slow response trials did not differ; see Methods section). Pupil size was significantly larger for fast trials compared to slow trials already more than 1 s before the gap onset (Fig. 3E).

In addition, we conducted a temporal response function analysis (Crosse et al., 2016) to ensure that the observed interaction of task difficulty and reward prospect is not resulting from a temporal shift in the pupil-size response due to response-time differences between conditions (see Fig. 2B). The time points of the noise onset, gap onset, and response time of the respective trial served as separate regressors to model the temporal response functions (TRF) to the different events. The TRF to the gap onset (Fig. 3G left) but not to the button response (Fig. 3G right) shows temporal variability between the different experimental conditions. However, importantly, we observed the interaction of task difficulty and reward for both the TRF to gap onset (1.82 s to 2.56 s) and to the button response (1.38 s to 2.22 s). Additionally, the pupil-size TRF was larger for hard compared to easy trials (TRF to gap onset: -2 s to -0.88 s and 1.36 s to 3.14 s; TRF to button response (0.96s to 2.54 s).

### Task difficulty, but not reward prospect, affects the auditory sensory response

We tested whether task difficulty and reward prospect influenced sensory responses, that is, responses in the auditory cortex. To this end, we focused on the M100 component in the 0.09 to 0.13 s time window after gap onset in temporal sensors. The M100 amplitude was larger for easy (i.e., longer gap duration) compared to hard trials (i.e., shorter gap duration; β = -.40, SE = .013, p = 1.33 × 10^-200^, log(BF) = 210.5; Fig. 4A & B). The effect for reward prospect (β = -.00, SE = .017, p > .9, log(BF) = -4.5), and the interaction were not significant (β = -.01, SE = .034, p > .9, log(BF) = -4.5).

### Task difficulty and reward prospect modulate the response-related evoked field

We also explored the late ERF component from 0.5 s onwards (Fig. 4A). Because the topography indicates the responses originate from the parietal cortex, the analysis of this component focused on parietal sensors. Sorting the gap-locked trials according to their response times (Fig. 4C) shows that this component is time-locked to the response rather than to the gap-onset (at 0 s in Fig. 4C), because the negative deflection succeeds the response time (dashed line). Hence, we analyzed this component time-locked to the button response (Fig. 4D) instead of time-locked to the gap-onset. Topographies for the time windows before, during, and after the button response are shown in Figure 4D & E, indicating that the response-related component is similar to the component we observed in the gap-locked analysis (Fig. 4C). The response-related component was stronger deflected for hard than easy trials (β = -.18, SE = .013, p = 1.54 × 10^-39^, log(BF) = 83), and for reward-relevant than reward-irrelevant trials (β = -.04, SE = .013, p = 1.33 × 10^-3^, log(BF) = 1; Fig. 4E). The interaction was not significant (β = -.05, SE = .026, p > .05, log(BF) = -2.5).

### Neither reward prospect nor task difficulty modulate pre-gap parietal alpha power

The time-frequency analysis shows that parietal alpha power in the 8-12 Hz frequency band increased over time after noise onset, up to when the gap occurred (Fig. 5C). However, alpha power in the 1.7 to 2.2 s time window after noise onset (i.e., 0.5 s before any gap could occur) showed no effect of task difficulty (β = .00, SE = .016, p > 0.9, log(BF) = -4.5), reward prospect (β = -.01, SE = .016, p > 0.9, log(BF) = -4.5) nor the interaction: β = -.02, SE = .032, p > 0.9, log(BF) = -4.5). Grouping trials according to hits and misses (irrespective of condition) showed the pre-gap alpha power increase for hits but not for miss trials. For hit trials, alpha power also decreased after gap onset (Fig. 5D).

Alpha power time courses were time-locked to gap onset for an analysis of the exact time window before gap onset. We investigated whether alpha power in the pre-gap time window (−0.5 to 0 s) was affected by our experimental manipulations. However, neither the effect of reward prospect (β = -.01, SE = .016, p > .5 , log(BF) = -4.5), task difficulty (β = -.02, SE = .016, p > .5, , log(BF) = -4), nor the interaction were significant (β = -.03, SE = .032, p > 0.5, , log(BF) = -4.5).

Although pre-gap alpha power was not modulated by task difficulty or reward prospect, we found a positive correlation between pre-gap alpha power and behavior. The probability of detecting a gap (for hard trials) was higher when pre-gap alpha power in parietal sensors was larger (Fig. 5E). This effect was present on the within-participant level (OR = 1.12, SE = .037, p = 7.81 × 10^-4^, log(BF) = 1.7), but not on the between-participants level (OR = .90, SE = .143, p > .5 , log(BF) = -1.6). There were no significant alpha-power effects on response time (within-participant: β = .03, SE = .029, p > 0.2, log(BF) = -3.5; between-participants: β = .19, SE = .142, p > 0.2, log(BF) = -1.1).

### Reward prospect modulates post-gap alpha-power suppression

We further explored the alpha-power suppression that followed the gap. Figure 6A suggests the alpha suppression is related to the gap rather than the button press, as illustrated by relation of the two dashed lines to the alpha-power suppression in Figure 6A. Task difficulty led to a post-gap alpha-power suppression that was only marginally significant for temporal sensors (β = -.04 SE = .018, p = 4.03 × 10^-2^, log(BF) = -2, Fig. 6C) and not significant for parietal sensors (β = -.03, SE = .018, p > .2 , log(BF) = -3.5, Fig. 6D). However, alpha power was more suppressed in reward-relevant compared to reward-irrelevant trials for temporal sensors (β = -.07, SE = .017, p = 3.33 × 10^-4^, log(BF) = 3) and parietal sensors (β = -.05, SE = .018, p = 1.53 × 10^-2^, log(BF) = -.5). For neither of the two sensor clusters was the interaction significant (temporal sensors: β = -.06, SE = .035, p > .1 , log(BF) = -3; parietal sensors: β = -.03, SE = .036, p > .4 , log(BF) = -4). Projecting gap-locked alpha power data into source space revealed qualitatively similar results to the results in sensor space, although not all effects reached statistical significance in source space (see Fig. 7).

**Figure 6.**
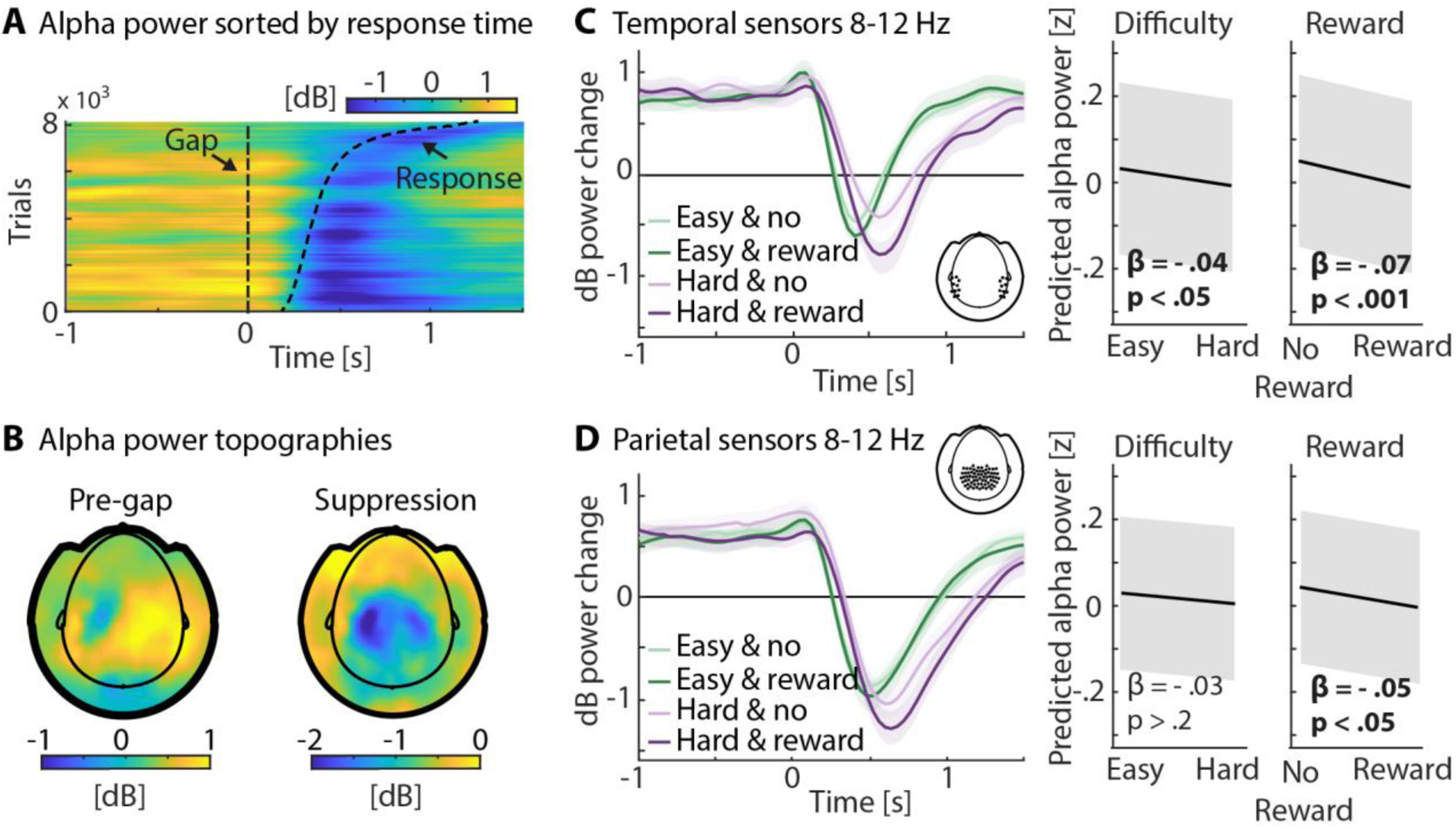
Post-gap alpha power analysis. **A:** Gap-locked alpha power trials sorted by response time. Data is averaged across parietal sensors see inset in D. Black lines represent the gap onset and the response time. Alpha-power suppression is rather time-locked to the gap-onset than to the response time. **B:** Left: Grand average topography for -.5 to 0s time-locked to gap onset. Right: Grand average topography for maximal alpha-power suppression after gap onset. **C:** Temporal sensors. Left: Alpha power averaged per condition across participants and temporal sensors (see inset). Error bands reflect the within-participants error. Right: linear mixed-effect model results of alpha-power suppression. Alpha-power suppression was larger for hard compared to easy and for reward-relevant compared to reward-irrelevant trials. **D:** Parietal sensors. Left: Alpha power averaged per condition across participants and parietal sensors (see inset). Error bands reflect the within-participants error. Right: linear mixed-effect model results of alpha-power suppression. Alpha-power suppression was larger for reward-relevant compared to reward-irrelevant trials.

**Figure 7.**
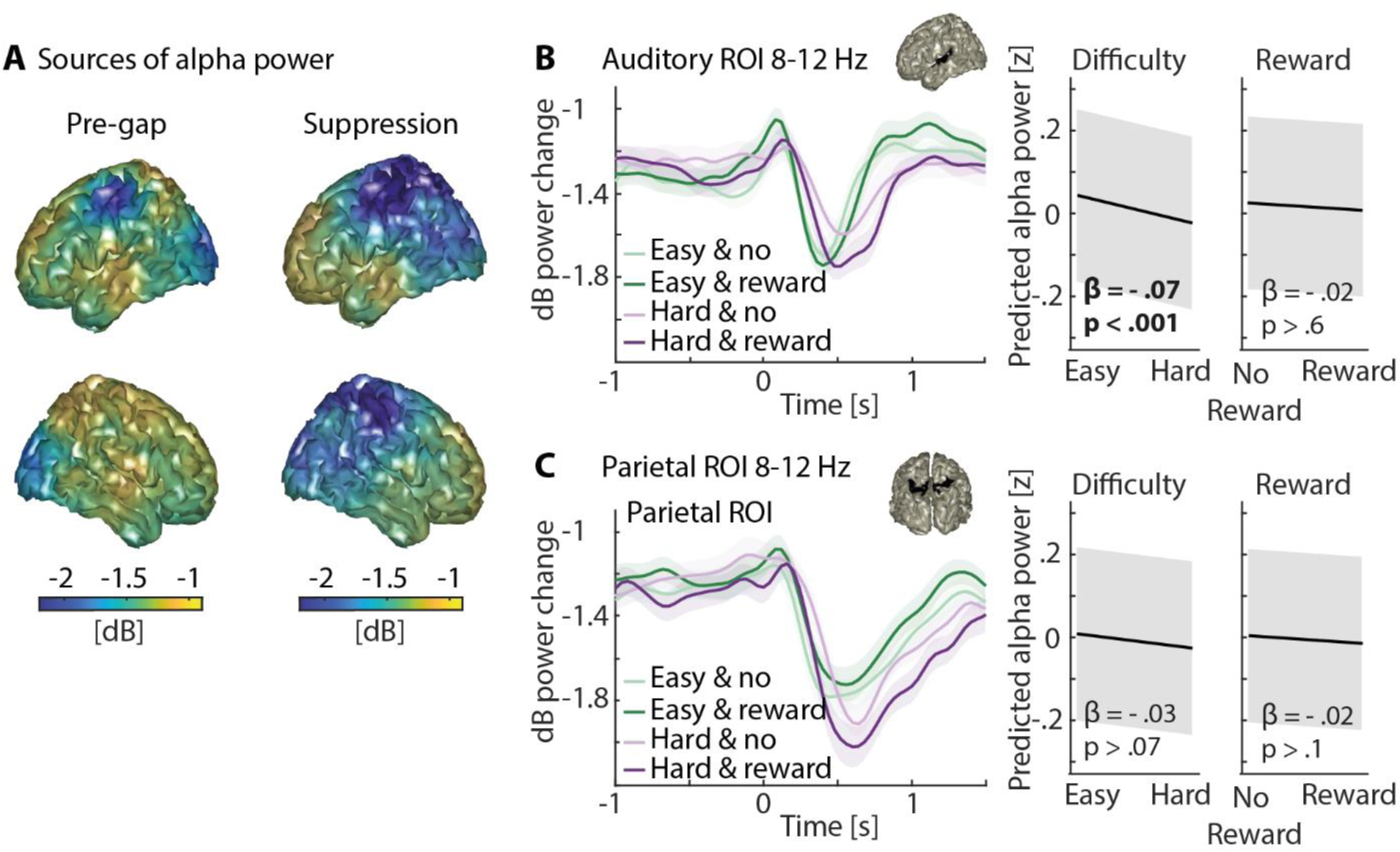
Source localized alpha power, time-locked to gap onset. **A:** Left: Grand average source plot for -.5 to 0 s time-locked to gap onset. Right: Grand average source plot for maximal alpha-power suppression after gap onset. **B:** Auditory ROI. Left: Source-projected alpha power for each condition. Inset shows the auditory ROI. Error bands reflect the within-participants error. Right: Results from a linear mixed-effect model predicting alpha-power suppression. Alpha-power suppression was larger for hard compared to easy trials (β = -.07, SE = .018, p = 9.64 × 10^-4^, log(BF) = 2.5). The effect of reward (β = -.02, SE = .018, p > .6, log(BF) = -4) and the difficulty × reward interaction (β = .02, SE = .036, p > .7, log(BF) = -4.5) were not significant. Direction of the effects is same as in sensor space. **C:** Same as in panel B for a parietal ROI. No significant effect was found (task difficulty: β = -.03, SE = 0.016, p > .7, log(BF) = -2.5; reward prospect: β = -.02, SE = .016, p > .1, log(BF) = -3.5; interaction: β = -.06, SE = .031, p > .1, log(BF) = -3) Direction of the effects is the same as in sensor space.

Correlating the M100-amplitude with the post-gap alpha-power suppression revealed a positive relationship. When the M100-amplitude across temporal sensors was enhanced, subsequent alpha power was less suppressed in both temporal (within-participant: β = .11, SE = .020, p = 2.74 × 10^-8^, log(BF) = 108; between-participants: β = .37, SE = .049, p = 6.26 × 10^-13^, log(BF) = 12.8) and parietal sensors (within-participant: β = .09, SE = .016, p = 2.74 × 10^-8^, log(BF) = 47; between-participants: β = .20, SE = .076, p = 1.36 × 10^-2^, log(BF) = 1.2).

### Larger pre-gap pupil size covaries with lower pre-gap parietal alpha power

To investigate the relationship between pupil size and brain measures, we calculated different linear mixed-effect models (Fig. 8). First, we tested whether the pre-gap pupil size is associated with auditory cortex responses to the gap (M100 amplitude; averaged across temporal sensors). There was no within-participant effect (β = -.01, SE = .018, p > .6, log(BF) = -6) nor a between-participants effect (β = -.07, SE = .053, p > .4, log(BF) = -1.73).

**Figure 8.**
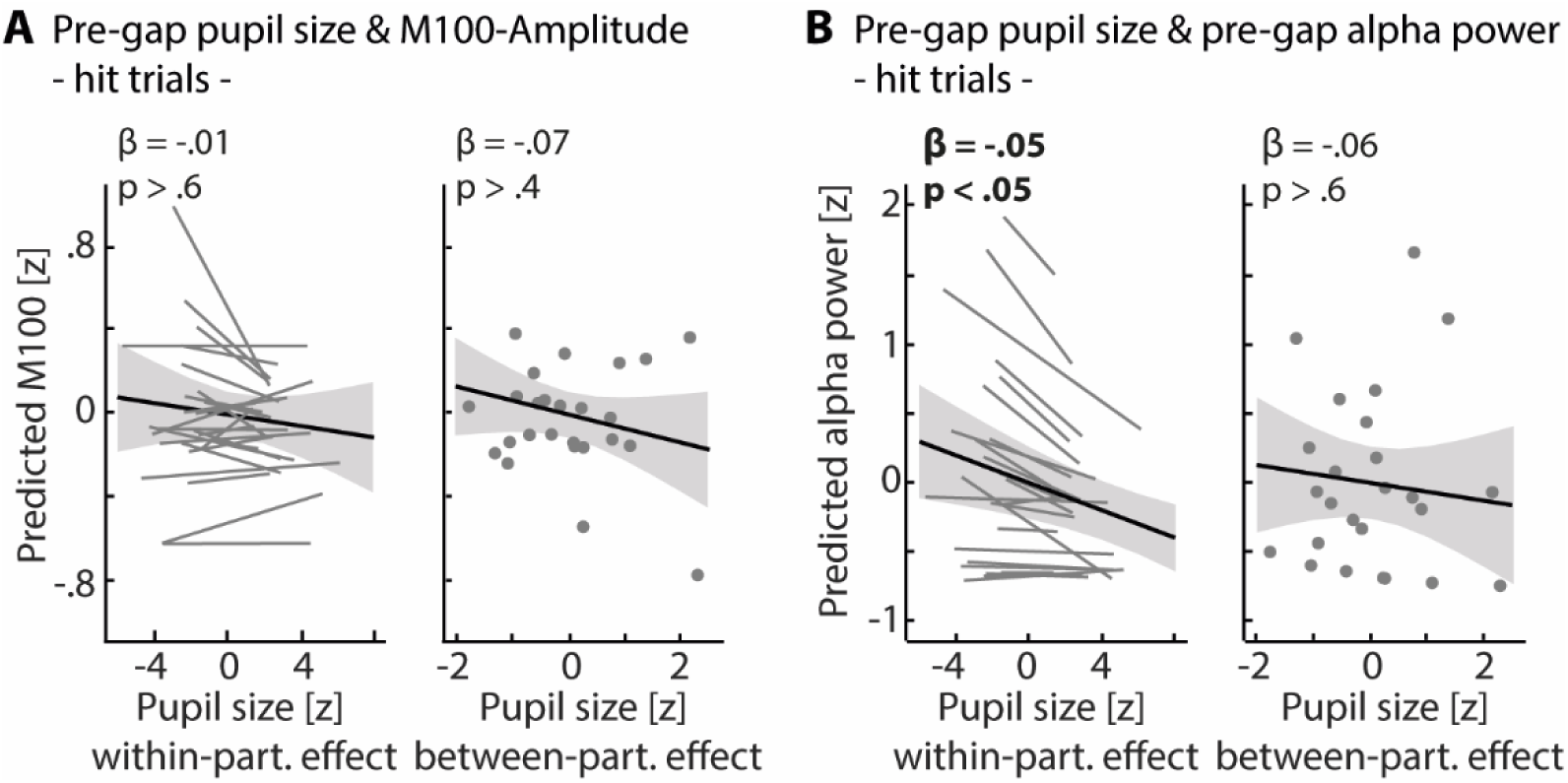
Relation between pupil size and brain measures. **A:** Relation between pre-gap pupil size and M100-amplitude. Linear mixed-effect modeling controlled for experimental conditions and was done on hit trials only. Pupil size was averaged across 0.5 s before gap-onset and baseline-corrected to the baseline window before the trial. M100 data was averaged across temporal sensors. No within-participant nor between-participants effect of pupil size on the M100-amplitude was found. **B:** Relation between pre-gap pupil size and pre-gap parietal alpha power. Linear mixed-effect modeling controlled for experimental conditions and was done on hit trials only. Pupil size was averaged across 0.5 s before gap-onset and baseline-corrected to the baseline window before the trial. Alpha power data was averaged across the same 0.5 s time window and across parietal sensors. A significant negative relationship between pupil size and alpha power was found on a within-participant level.

Second, we investigated whether the pre-gap pupil size is associated with pre-gap parietal alpha power (averaged across parietal sensors). A larger pupil size was associated with lower parietal alpha power on a within-participant level (β = -.05, SE = 0.02, p = 1.25 × 10^-2^, log(BF) = 5.5; Fig. 8 B). There was no significant effect on a between-participants-level (β = -.06, SE = .104, p > .6, log(BF) = -1.64). Importantly, controlling for possible changes of alpha power over time via including the trial number across the experiment as an additional regressor into the model had no effect on the results.

## Discussion

In the current study, we investigated a listener’s motivation affecting behavior, pupil size, and neural activity during a challenging auditory detection task. Participants performed better, and pupil size increased when listeners were more motivated compared to when they were less motivated by reward prospects. Neural oscillatory activity in parietal regions increased throughout the presentation of the noise sound, but this power increase was independent of task difficulty and reward prospect. Auditory sensory responses to auditory target stimuli were not modulated by reward prospect. However, both task difficulty and reward prospect independently enhanced post-gap activity. In sum, pupil-size changes index cognitive investment during the interplay of listening demand and motivation, whereas motivation impacts neural indices post-stimulus rather than preparatory attentional modulation. The current study thus shows the complex and distinct impacts of motivation and listening demands on different neurophysiological systems.

### Motivational state moderates the effect of listening demand on pupil size

Pupil size increased as listening demands increased, as expected (Kadem et al., 2020; Kahneman & Beatty, 1966; Koelewijn et al., 2012; Kraus et al., 2023b; Ohlenforst et al., 2018; Wendt et al., 2016; Winn et al., 2015; Zekveld et al., 2010). Previous work has also shown that the pupil size is sensitive to individuals giving up listening under impossible listening conditions (Herrmann & Ryan, 2024; Ohlenforst et al., 2017; Zekveld & Kramer, 2014). The observation of the current study that a low motivational state is associated with a smaller pupil size, especially under high listening demands (see Fig. 3B), is consistent with the literature on ‘giving up’ listening. People may invest less cognitively when they are little motivated (Brehm & Self, 1989; Herrmann & Johnsrude, 2020; Koelewijn et al., 2018; Kraus et al., 2023a; Pichora-Fuller et al., 2016). A consideration in the current study may be that response times differed between conditions and that response time temporally aligned with the pupil dilation. However, the moderating effect of the motivational state on the task difficulty effect on pupil size remains qualitatively the same when controlling for the potential confound of the different response times (see Fig. 3 G).

Critically, the behavioral results by themselves suggest that the influence of motivation is independent of the influence of listening demand (two main effects; Fig. 2). However, the pupil-size results indicate, consistently with cognitive control frameworks (Brehm & Self, 1989; Parro et al., 2018; Yee & Braver, 2018), that motivation is most impactful under difficult compared to easy listening demands. Consequently, based on motivational intensity theory (Brehm & Self, 1989; Richter, 2016) the individual motivational state is another important determinant of the amount of effort allocated during listening besides the level of listening demand. The here and previously found increases in pupil size and response speed with heightened motivation (Kraus et al., 2023a) highlights the importance of taking listeners’ motivation into account when assessing their invested effort. Our pupil size data thus provide an important confirmation to examine motivational impacts on the neural level.

### Expectations about different levels of listening demand are not sufficient to modulate alpha power

Parietal alpha power increased throughout the presentation of the noise up to when individuals detected the gap (Fig. 5C). Alpha power thus showed sensitivity to when in time attention is allocated, followed by a post-stimulus suppression (Herrmann et al., 2023). However, in contrast to previous work (Obleser et al., 2012; Petersen et al., 2015; Wisniewski et al., 2017; Wöstmann et al., 2015) and our hypothesis, we did not observe an increase in parietal alpha power with increasing task demand nor when listeners could expect a reward for performing well. One important difference between our and these former studies is that different tasks and time windows of interest were employed. Most of the studies that report demand-related effects on parietal alpha power during listening observed them in a memory task, during the retention period when participants held a stimulus in memory to compare it with a later probe stimulus (Obleser et al., 2012; Petersen et al., 2015; Wisniewski et al., 2017; Wöstmann et al., 2015). In the present study, we investigated alpha power changes in a period – leading up to a behaviorally relevant auditory event – during which our experimental conditions were acoustically identical (as opposed to some of the previous studies). It thus appears that top-down knowledge about the difficulty of an upcoming event is insufficient to drive changes in parietal alpha power (but also see Herrmann et al. (2023), indicating additional complexities). Palva & Palva (2011) suggest that distinguishing between endogenous vs. exogenous task demands is critical to understanding alpha-power dynamics. Manipulation of task demand via the saliency of a stimulus is an exogenous manipulation (Obleser et al., 2012; Petersen et al., 2015; Wisniewski et al., 2017; Wöstmann et al., 2015), whereas adjusting expectations about demand level is an endogenous manipulation (present study). Therefore, we suggest that changes in alpha power due to listening demand may only be observed under specific task and stimulus conditions but do not generally vary with different levels of cognitive resource recruitment during listening.

### Not auditory sensory processing but post-stimulus alpha power is modulated by motivational state

Auditory sensory responses (M100 amplitude) were enhanced for a long compared to a short gap duration – and thus for the more salient target – but not for variations in motivation. The M100 and the N100, the electric equivalent, are known to respond most strongly to changes in physical stimulus properties (Frank et al., 2020; Hansen & Hillyard, 1980; Näätänen & Picton, 1987; Paiva et al., 2016). Nevertheless, attentional modulations of the M100/N100 have been observed previously but often limited to spatial tasks (Ding & Simon, 2012; Hillyard et al., 1973; Kraus et al., 2021; O’Sullivan et al., 2015; Orf et al., 2023; Woldorff & Hillyard, 1991). It appears, however, that the attentional boost through motivation is insufficient to affect auditory sensory responses.

In contrast to the auditory sensory processing, we observed a response-related neural signal that scaled with task difficulty and reward prospect. Research on the impact of reward on stimulus processing often focuses on the feedback-related negativity (FRN). The FRN is a component that peaks around 200 ms after the feedback about an earned or lost reward (Glazer et al., 2018; Holroyd & Coles, 2002). The FRN is thought to scale with the degree to which the actual feedback deviates from the expected feedback (Frömer et al., 2021; Sambrook & Goslin, 2015) and is modulated by reward expectation (M. X. Cohen et al., 2007). In the present study, participants knew whether they were in a reward-irrelevant or a reward-relevant trial and how difficult target detection was. Moreover, the response participants made upon gap detection was the only feedback that they received about possibly gaining a reward. Detecting a gap in hard trials may have led to a more unexpected feedback signal (i.e., positive surprise) than in easy trials because participants knew that it would be less likely for them to detect the gap in hard trials. This feedback signal may be greater in the reward-relevant than the reward-irrelevant condition. We, therefore, interpret the observed effect of motivation as a positive surprise related to the detection of the gap.

Although there was no change in pre-gap alpha power nor in auditory sensory processing (M100) due to changes in motivational state, the post-gap alpha-power suppression was stronger during high compared to low motivation, independently of task demand. The suppression of alpha power is thought to be a sign of the facilitation, and therefore gating, of the processing of sensory input (Jensen & Mazaheri, 2010; Klimesch et al., 2007; Palva & Palva, 2011; Pfurtscheller, 2003). In visual attention tasks, reward prospect enhanced post-target encoding (Hall-McMaster et al., 2019) and neural processing in attention networks (Small et al., 2005), suggesting motivation influences of attentional top-down mechanisms on sensory processing. Both the response-related neural signal and the post-stimulus alpha-power suppression suggest motivation affects top-down mechanisms during effortful listening.

### Discrepancy between changes in pupil size and alpha power

Both parietal alpha power and pupil size increased over the trial time course toward the gap, which proved beneficial for behavior. However, the degree to which both measures were affected by our experimental manipulations differed. Pupil size was greater for higher task demands and modulated by motivation, whereas parietal alpha power prior to the gap did not change for either manipulation. Research investigating the relations between both metrics is rare. The few studies that observed simultaneous demand-driven dynamics in pupil size and alpha power manipulated the demand via the stimulus itself, for example, by adding different levels of background noise to speech or using noise-vocoded speech at different levels (Ala et al., 2020; McMahon et al., 2016; Miles et al.,2017). As outlined above, the present study had no acoustic differences in the pre-gap analysis window, which may explain the absence of alpha-power changes. In a recent study using the same auditory paradigm in an audiovisual dual-task, we also found that changes driven by listening demands were more prominent in pupil size than in parietal alpha power (Kraus et al., 2023b).

The current study also showed that pre-gap pupil size and pre-gap parietal alpha power are negatively related (see also Kraus et al., 2023b). Although pupil size and alpha power were differently sensitive to listening demand and motivation, a common underlying mechanism seems to affect both pupil size and parietal alpha power but in opposite directions. Noradrenergic dynamics originating from the locus coeruleus may be one candidate. Pupil size variations are correlated with activity in the locus coeruleus (Aston-Jones & Cohen, 2005b), and the pupil dilates in response to LC-stimulation (Liu et al., 2017). Increased noradrenergic activity emerging from LC is linked to optimizing task performance and task engagement (Aston-Jones & Cohen, 2005a). The noradrenergic influence on thalamic modulations (McCormick, 1998) and the thalamic connection to cortical oscillatory activity (Steriade, 2000) provide evidence for a neural mechanism affecting both pupil size and neural oscillatory alpha power (Dahl et al., 2022).

## Conclusion

The present study confirms that pupil size robustly indexes cognitive investment during listening and provides an integrated read-out of an individual’s motivational state. However, the neural mechanisms that encode listening demand and motivational state reveal a more nuanced picture: Expectation of a more demanding or more important (i.e., reward-relevant) sensory input does not suffice to elicit variations in neural alpha power. Auditory sensory processing was also not affected by a listener’s motivational state. Importantly, we observed that the known alpha-power suppression in the wake of an auditory target event appears amplified under top-down motivational regulation. In sum, while pupil size poses an integrated read-out of listening demand and motivation, both dimensions selectively affect sensory and attentional post-processing aspects of auditory neurophysiology.

## Acknowledgments

We thank Gloria Lai and Tazeen Atif for their assistance with data collection. This work was supported by Deutsche Forschungsgemeinschaft (DFG; grant number HE 7857/1-1) awarded to BH. BH is supported by the Natural Sciences and Engineering Research Council of Canada (Discovery Grant: RGPIN-2021-02602) and the Canada Research Chair program (CRC-2019-00156).

## Author contributions

F.K., B.H. and J.O. designed research, F.K. performed research, B.R. provided resources, F.K., J.O., and B.H. analyzed data, F.K. visualized the data, F.K. wrote first draft of the manuscript, F.K., B.R., B.H., and J.O. edited the manuscript.

